# High-throughput MicroED for probing ion channel dynamics

**DOI:** 10.1101/2025.02.19.639122

**Authors:** Marc J Gallenito, Max TB Clabbers, Jieye Lin, Johan Hattne, Tamir Gonen

## Abstract

Ion channels play a crucial role in ion transport and are integral to fundamental physiological processes. Therefore, understanding channel structures is essential for elucidating the mechanisms of ion permeation and selectivity beyond what can be predicted by computational simulations. Visualizing dynamics at high resolution, however, remains a significant challenge by structural techniques. In this study, we apply high-throughput Microcrystal Electron Diffraction (MicroED) to explore the structural dynamics of two ion channels, the non-selective ion channel NaK and its mutant, NaK2CNG. This approach utilizes automated data collection and processing to capture distinct structural substates from a large number of microcrystals, offering a deeper understanding of ion channel mechanisms. From a subset of NaK structures, we observed consistent sodium binding at specific sites. In contrast, NaK2CNG appears more dynamic and undergoes dilation of the selectivity filter upon potassium binding. Further, the conduction state of NaK2CNG appears to be influenced by channel gating. Comparative analysis of these structures reveals that non-selectivity arises from the plasticity of the selectivity filter, allowing dynamic control over ion passage. These studies, demonstrate the potential to employ high-throughput MicroED as a technique to address persistent questions regarding ion channel permeation, complementing current computational molecular dynamics studies. We anticipate that this approach will enhance future computational models, leading to more accurate predictions of ion channel behavior and providing a more comprehensive view of transport dynamics.

## Introduction

Characterization of ion channel permeation and selectivity is fundamental to our understanding of biophysics and physiology, as these play key roles in essential biological processes. The transport of ions, whether selective or not, plays a critical role in maintaining homeostasis, facilitating cellular functions, and enabling cell-cell communication^1^. Disruption in ion channel function, often due to loss-of-function mutations, can lead to a range of deleterious phenotypes, including cardiac arrhythmias, neurological disorders, and metabolic conditions^2,3^. Understanding the molecular mechanisms that govern these processes is, therefore, crucial for addressing these channelopathies. Dissecting the molecular mechanisms of ion channels requires a detailed exploration of how specific mutations affect ion conduction and selectivity, and examination of the protein structural dynamics at play during these processes. Unraveling the complexities of ion channels provide insights to enhance our understanding of cellular physiology and inform therapeutic strategies.

Early structural studies using the prototypical K^+^-selective channel KcsA revealed that the selectivity filter is encircled by carbonyl oxygens of the peptide backbone in the “signature sequence” Val-Gly-Tyr-Gly-Asp^4,5^. The structures of KcsA support the “soft knock-on” mechanism, in which K^+^ ions and water molecules are transported in a coordinated fashion by alternating through the selectivity filter of the channel^6,7^. This mechanism became the founding principle for ion conduction models and provided the framework for computational and experimental method for over the following decade. The alternating ion-water transport mechanism stood as the sole mechanism for ion transport until advancements in computational electrophysiology introduced the model that preclude water intervention, permitting ion-ion contacts. This model is termed the “direct knock-on” mechanism^8,9^.

NaK is a non-selective ion channel^10^. The overall organization of NaK is similar to that of KcsA, except for the selectivity filter, which allows the channel to be permeable to Na^+^, K^+^, and other divalent cations^10,11^. Initial X-ray structures of NaK showed a rigid selectivity filter^10,12^. Recent NMR and molecular dynamics simulations, however, suggested that a dynamic selectivity filter is essential for its non-selectivity ^13^. Consequently, the mechanisms underlying the non-selectivity of NaK channels remain to be determined. Engineering of the NaK selectivity filter generated two variants, NaK2CNG and NaK2K, demonstrating that small modifications result in significant changes in ion selectivity and conduction^14,15^. NaK2CNG, which mimics eukaryotic cyclic nucleotide-gated (CNG) channels, has been used as a model for studying non-selective systems. Although NaK2CNG and K^+^-selective channels share similarities in the number of oxygen-lined sites in their selectivity filters, they exhibit distinct permeation preferences^15,16^.

Ion channels are some of the fastest-acting proteins in nature, with a transport rate of several million ions per second^17^. For this reason, capturing the dynamic nature of ion conduction poses significant challenges for current structural methods, including x-ray crystallography and single particle analysis (SPA) cryo-EM. While SPA has provided insights into selectivity filter dynamics and channel activation, determining ion occupancy—especially for mobile ions within the selectivity filter—remains challenging^18,19^. Moreover, SPA is also limited to rather large proteins and many ion channels are too small for studying by this method. Conversely, although X-ray structures have contributed to computational studies, this method necessitates large protein crystals, which typically limits the number of attainable structures often modeling only a single conformation/state. For these reasons, channel dynamics is traditionally probed by in silico methods like molecular dynamics simulations, Brownian dynamics, and quantum-mechanical simulations^20–27^. These methods are used to model real-time ion movements under simulated physiological conditions.

The cryo-EM method microcrystal electron diffraction (MicroED) offers an alternative strategy to probe dynamics using vanishingly small crystals ^28^. Here, crystals that are a billionth the size which is needed for X-ray diffraction are used in an electron microscope and data are collected under cryogenic conditions. MicroED has allowed structure determination of difficult proteins and unambiguous assignment of cation and amino acid charges^29,30^. The MicroED structure of NaK was one of the first membrane protein structures determined by this technique and demonstrate that distinct ion channel states can be captured even using protein in crystal lattices ^29^

Here, we utilized high-throughput MicroED, originally reported for analyzing small molecule mixtures and compositional studies ^31^, to investigate the dynamics of the NaK channel and its mutant NaK2CNG in protein crystal form. We demonstrate that this approach is viable even in the context of a crystal because as channel permeability occurs at the selectivity filter, which is far removed from crystal lattice contacts in these ion channels. We selected NaK due to its ability to form flat crystals that are amendable for MicroED diffraction without further physical manipulation such as FIB milling^29,32^. Additionally, the crystal symmetry is sufficiently high to allow for data acquisition for complete structure determination from individual crystals, eliminating the need for averaging across multiple crystals. Finally, the axis of symmetry of these crystals does not overlap with the axis of the selectivity filter so charge densities could be identified unambiguously. We adapted the high-throughput procedures for protein structure data collection and determination, enabling us to systematically probe hundreds of ion channel crystals and multiple substates rather than relying on a single static model.

## Results and Discussion

### High-throughput MicroED

NaK and NaK2CNG microcrystals were grown in Tris pH 8.0 and 60-80% (±)-2-methyl-2,4-pentadiol (MPD). Consistent with past studies, crystals were generally plate-like with varying x and y dimensions, and a thickness suitable for electron diffraction of less than 0.5 µm^29^. Figure 1 illustrates the general workflow for high-throughput MicroED. Full grid montages at low magnification were collected to identify intact squares with thin ice. To aid in crystal picking, medium-magnification images of grid squares were collected. Objects that resemble a crystal, those with sharp edges, square or rectangular shapes, were manually selected, wherein points for data collection were placed. For larger crystals, multiple points were placed within each crystal for initial diffraction tests. For each position, the crystal quality was evaluated by collecting a single diffraction pattern without rotation. Initial diffraction test takes less than 2s per point and only exposes the crystal to a minimal amount of electron fluence (<0.002 e^-^/Å^2^), conferring negligible radiation damage^33^. Crystals of low-quality or no diffraction were eliminated from full data collection. Continuous rotation MicroED data were collected from crystals showing strong preliminary diffraction to high resolution. A total of 75 degrees worth of continuous rotation data were collected from each crystal using total fluence of less than 0.84 e^-^/Å^2 34^.

**Figure 1.**
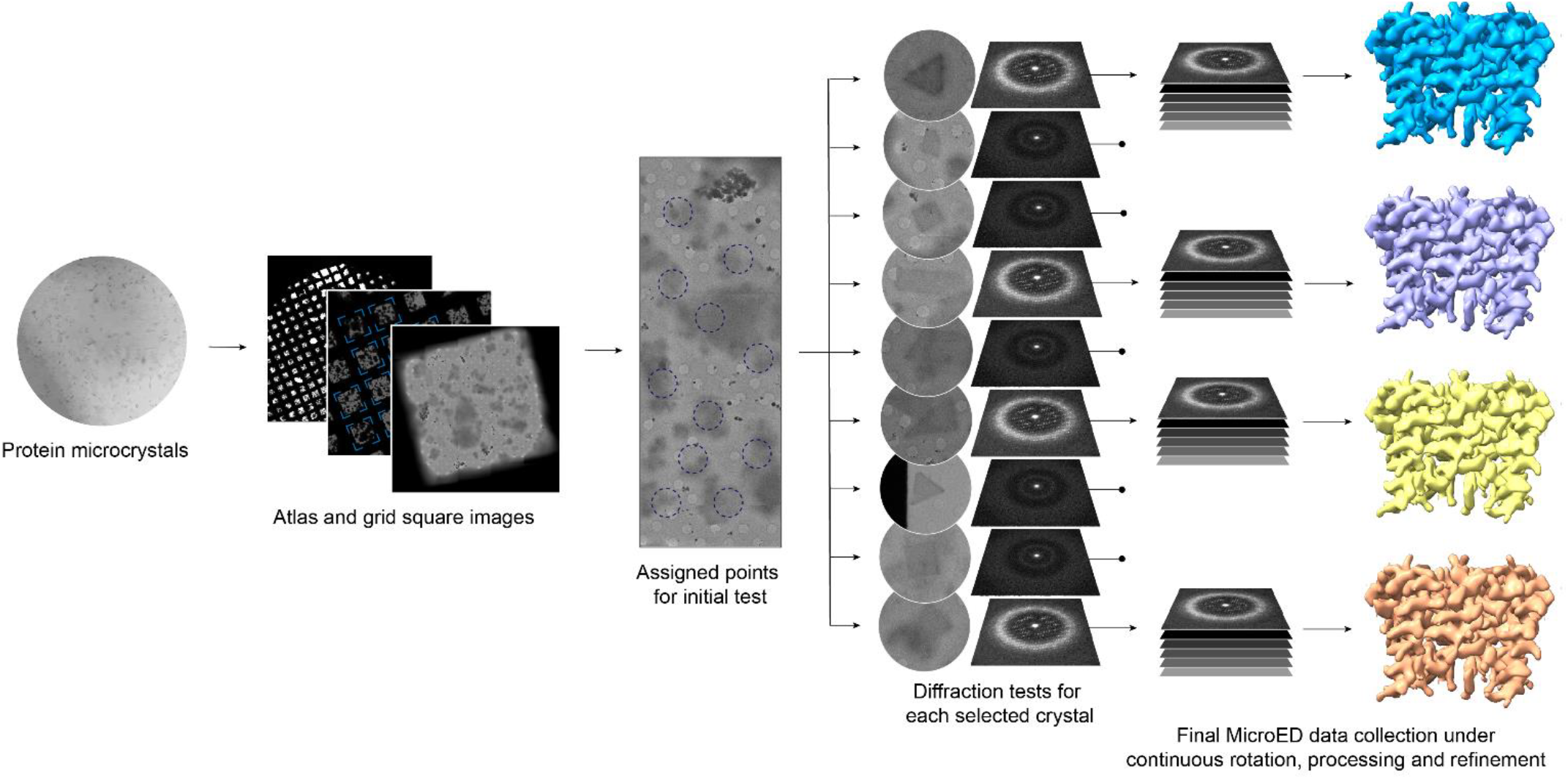
Schematic flow of high-throughput MicroED.

Plate-like crystals tend to disperse on TEM grids with a preferred orientation. Therefore, MicroED data were collected using an optimized angle strategy to cover a larger rotation range and to take advantage of the high crystal symmetry of NaK for high data completeness. Using this approach each crystal could typically yield >80% data independently without the need to merge data from multiple crystals (Table 1). Since NaK2CNG produces similar crystal morphology under the same conditions as NaK, no further modifications in the method were needed for MicroED data collection. While we diffracted from hundreds of crystals, only a handful yielded high resolution data with high completeness. This is typically observed in membrane protein crystallography where crystal quality varies greatly even within a crystallization condition. After culling, we systematically chose 10 individual data sets for final structure solution: 6 structures of NaK and 4 for NaK2CNG with the best structural parameters circumventing the need for merging datasets across multiple crystals (Table 1).

**Table 1.** MicroED data collection and refinement statistics for NaK and NaK2CNG.

|  | NaK 1 | NaK 2 | NaK 3 | NaK 4 | NaK 5 | NaK 6 |
| --- | --- | --- | --- | --- | --- | --- |
| **Resolution range** | 34.1 - 2.8 (34.1 - 2.8) | 34.13 - 2.9 (34.13 - 2.9) | 32.95 - 2.8 (3.53 - 2.8) | 34.12 - 3.0 (34.12 - 3.0) | 54.46 - 2.8 (54.46 - 2.8) | 34.38 - 3.0 (34.38 - 3.0) |
| **Space group** | I 4 | I 4 | I 4 | I 4 | I 4 | I 4 |
| **Unit cell** | 68.197 68.197 90.451 90 90 90 | 68.253 68.253 92.374 90 90 90 | 68.428 68.428 89.982 90 90 90 | 68.242 68.242 91.179 90 90 90 | 68.338 68.338 90.162 90 90 90 | 68.753 68.753 89.972 90 90 90 |
| **Total reflections** | 20865 (20865) | 21297 (21297) | 20068 (9614) | 10174 (10174) | 20243 (20243) | 17713 (17713) |
| **Unique reflections** | 11831 (11831) | 11573 (11573) | 11477 (5510) | 6443 (6443) | 10842 (10842) | 10490 (10490) |
| **Multiplicity** | 1.8 (1.8) | 1.8 (1.8) | 1.7 (1.7) | 1.6 (1.6) | 1.9 (1.9) | 1.7 (1.7) |
| **Completeness (%)** | 74.67 (74.67) | 80.52 (80.52) | 82.41 (83.31) | 65.03 (65.03) | 74.96 (74.96) | 74.90 (74.90) |
| **Mean I/sigma(I)** | 1.45 (1.45) | 1.58 (1.58) | 1.44 (0.56) | 2.49 (2.49) | 1.97 (1.97) | 1.29 (1.29) |
| **Wilson B-factor** | 56.18 | 57.55 | 57.95 | 66.62 | 56.36 | 65.08 |
| **R-merge** | 0.2345 (0.2345) | 0.2839 (0.2839) | 0.2327 (0.7258) | 0.1481 (0.1481) | 0.2383 (0.2383) | 0.3314 (0.3314) |
| **R-meas** | 0.3055 (0.3055) | 0.3708 (0.3708) | 0.3072 (0.9598) | 0.1928 (0.1928) | 0.3053 (0.3053) | 0.4384 (0.4384) |
| **R-pim** | 0.1929 (0.1929) | 0.2354 (0.2354) | 0.1979 (0.6201) | 0.1216 (0.1216) | 0.1879 (0.1879) | 0.2833 (0.2833) |
| **CC1/2** | 0.966 (0.966) | 0.955 (0.955) | 0.969 (0.138) | 0.979 (0.979) | 0.977 (0.977) | 0.95 (0.95) |
| **CC\*** | 0.991 (0.991) | 0.988 (0.988) | 0.992 (0.492) | 0.995 (0.995) | 0.994 (0.994) | 0.987 (0.987) |
| **Reflections used in refinement** | 3845 (3845) | 3812 (3812) | 4244 (2131) | 2741 (2741) | 3862 (3862) | 3166 (3166) |
| **Reflections used for R-free** | 155 (155) | 205 (205) | 225 (103) | 145 (145) | 204 (204) | 139 (139) |
| **R-work** | 0.2535 (0.2535) | 0.2748 (0.2748) | 0.3098 (0.3642) | 0.2486 (0.2486) | 0.2517 (0.2517) | 0.2502 (0.2502) |
| **R-free** | 0.3092 (0.3092) | 0.3083 (0.3083) | 0.3487 (0.3977) | 0.2841 (0.2841) | 0.3062 (0.3062) | 0.2836 (0.2836) |
| **Number of non-hydrogen atoms** | 1311 | 1424 | 1297 | 1425 | 1424 | 1426 |
| **macromolecules** | 1295 | 1418 | 1292 | 1418 | 1418 | 1418 |
| **ligands** | 13 | 6 | 5 | 7 | 6 | 8 |
| **solvent** | 3 | 0 | 0 | 0 | 0 | 0 |
| **Protein residues** | 165 | 182 | 165 | 182 | 182 | 182 |
| **RMS(bonds)** | 0.001 | 0.001 | 0.001 | 0.001 | 0.002 | 0.002 |
| **RMS(angles)** | 0.32 | 0.43 | 0.35 | 0.39 | 0.47 | 0.42 |
| **Ramachandran favored (%)** | 93.79 | 96.07 | 99.38 | 96.07 | 97.19 | 97.75 |
| **Ramachandran allowed (%)** | 6.21 | 3.93 | 0.62 | 3.93 | 2.81 | 2.25 |
| **Ramachandran outliers (%)** | 0 | 0 | 0 | 0 | 0 | 0 |
| **Rotamer outliers (%)** | 0 | 1.23 | 0 | 0 | 0 | 0 |
| **Clashscore** | 6.81 | 2.76 | 1.9 | 2.76 | 1.73 | 1.72 |
| **Average B-factor** | 62.98 | 53.99 | 68.07 | 69.68 | 67.48 | 69.93 |
| **macromolecules** | 62.89 | 53.99 | 68.14 | 69.71 | 67.54 | 70.05 |
| **ligands** | 79.37 | 53.85 | 51.43 | 62.08 | 52.52 | 48.49 |

|  | NaK2CNG 1 | NaK2CNG 2 | NaK2CNG 3 | NaK2CNG 4 |
| --- | --- | --- | --- | --- |
| **Resolution range** | 34.56 - 2.8 (3.21 - 2.8) | 33.58 - 2.8 (3.52 - 2.8) | 34.52 - 2.8 (34.52 - 2.8) | 49.08 - 2.8 (3.53 - 2.8) |
| **Space group** | I 4 | I 4 | I 4 | I 4 |
| **Unit cell** | 69.117 69.117 92.236 90 90 90 | 69.234 69.234 92.286 90 90 90 | 69.04 69.04 92.098 90 90 90 | 69.414 69.414 91.999 90 90 90 |
| **Total reflections** | 21980 (7174) | 21528 (11127) | 18035 (18035) | 22019 (11104) |
| **Unique reflections** | 12181 (3984) | 12615 (6398) | 10317 (10317) | 12219 (6150) |
| **Multiplicity** | 1.8 (1.8) | 1.7 (1.7) | 1.7 (1.7) | 1.8 (1.8) |
| **Completeness (%)** | 80.90 (81.81) | 83.26 (84.31) | 71.58 (71.58) | 81.05 (81.91) |
| **Mean I/sigma(I)** | 1.76 (0.73) | 1.59 (0.58) | 1.56 (1.56) | 2.02 (0.69) |
| **Wilson B-factor** | 50.87 | 50.97 | 44.98 | 57.22 |
| **R-merge** | 0.2395 (0.6551) | 0.3094 (0.8334) | 0.2864 (0.2864) | 0.212 (0.6774) |
| **R-meas** | 0.3134 (0.8659) | 0.4181 (1.127) | 0.3726 (0.3726) | 0.2766 (0.887) |
| **R-pim** | 0.1991 (0.5585) | 0.2785 (0.7515) | 0.2347 (0.2347) | 0.1749 (0.5644) |
| **CC1/2** | 0.971 (0.0649) | 0.971 (0.145) | 0.964 (0.964) | 0.978 (0.216) |
| **CC\*** | 0.993 (0.349) | 0.993 (0.503) | 0.991 (0.991) | 0.994 (0.596) |
| **Reflections used in refinement** | 4333 (1448) | 4491 (2252) | 3830 (3830) | 4393 (2209) |
| **Reflections used for R-free** | 444 (149) | 236 (112) | 200 (200) | 232 (108) |
| **R-work** | 0.2626 (0.3200) | 0.2885 (0.3495) | 0.2737 (0.2737) | 0.2459 (0.3213) |
| **R-free** | 0.2925 (0.3278) | 0.3258 (0.4151) | 0.3375 (0.3375) | 0.2931 (0.3744) |
| **Number of non-hydrogen atoms** | 1410 | 1409 | 1412 | 1412 |
| **macromolecules** | 1402 | 1402 | 1402 | 1402 |
| **ligands** | 7 | 7 | 9 | 9 |
| **solvent** | 1 | 0 | 1 | 1 |
| **Protein residues** | 180 | 180 | 180 | 180 |
| **RMS(bonds)** | 0.002 | 0.001 | 0.002 | 0.002 |
| **RMS(angles)** | 0.47 | 0.39 | 0.48 | 0.41 |
| **Ramachandran favored (%)** | 96.59 | 98.3 | 95.45 | 98.86 |
| **Ramachandran allowed (%)** | 3.41 | 1.7 | 3.98 | 1.14 |
| **Ramachandran outliers (%)** | 0 | 0 | 0.57 | 0 |
| **Rotamer outliers (%)** | 0 | 0 | 0 | 0 |
| **Clashscore** | 2.08 | 2.08 | 3.12 | 1.04 |
| **Average B-factor** | 57.07 | 51.7 | 47.36 | 61.08 |
| **macromolecules** | 57.17 | 51.68 | 47.28 | 61.11 |
| **ligands** | 41.15 | 56.43 | 64.27 | 56.66 |
| **solvent** | 29.83 |  | 12.9 | 50.57 |

### Structures of NaK and NaK2CNG

NaK and NaK2CNG share the canonical Na^+^ channel morphology^10,11,15,16^. Within the *I*4 space group, each asymmetric unit contains two monomers arranged in a head-to-head fashion (Figure 2A). Individual monomers are ordered into two distinct tetramers, similar to previous studies^12,29^. Each monomer consists of two helices and contributes one loop at the 4-fold axis of the channel to form the selectivity filter (SF). The various NaK structures displayed difference map peaks within the SF that would suggest ion occupancy unique to each channel (Figure 2B). Indeed, after several rounds of refinement, the ions placed at different positions along the SF could be resolved, represented in Figure 2C.

**Figure 2.**
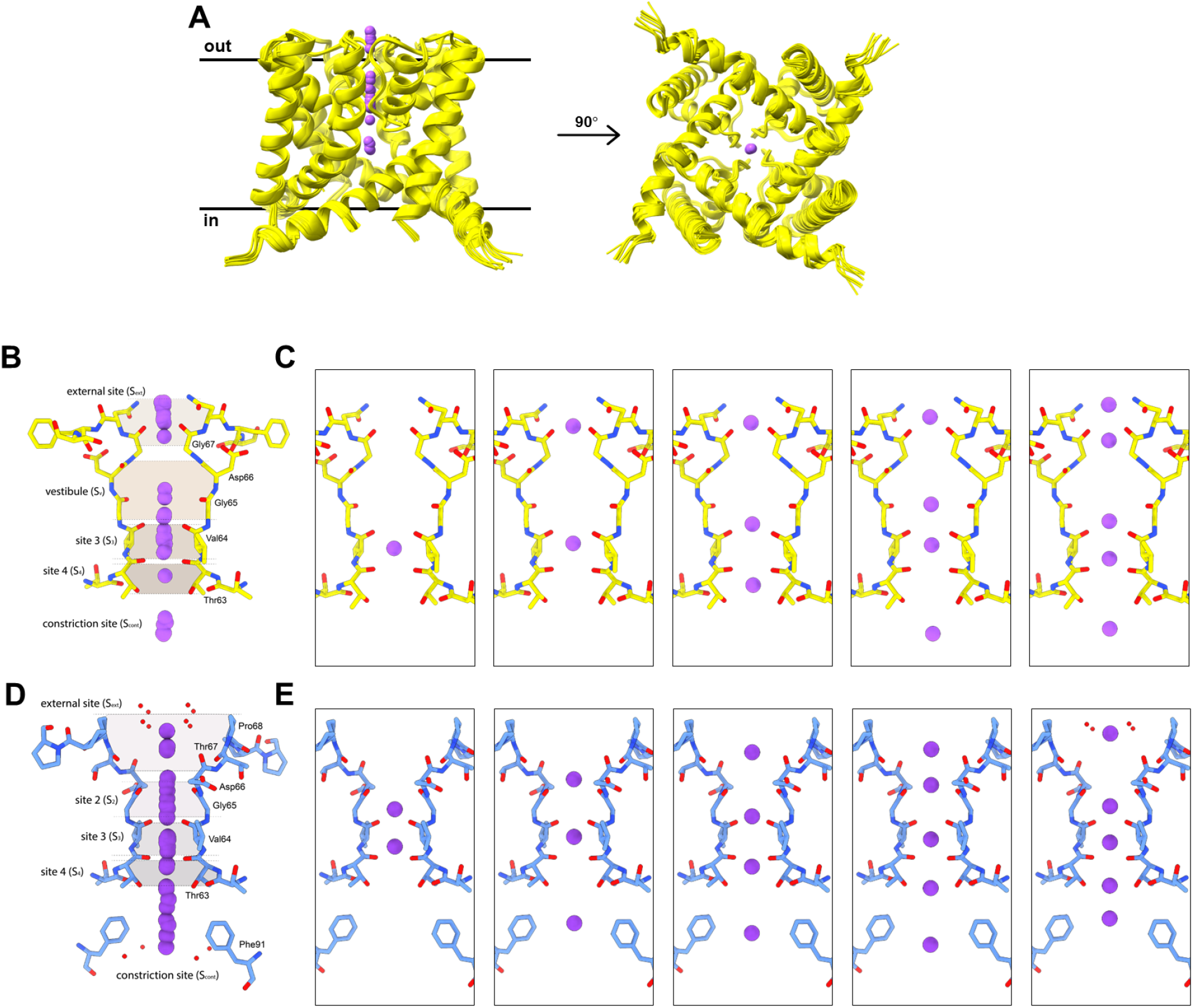
MicroED structures of NaK and NaK2CNG. (A) Alignment of tetrameric NaK structures. Sodium (B) and potassium (D) binding sites along the selectivity filter and representative sodium (C) and potassium (E) binding sites from all structures.

Structural alignment and analysis of all 12 structures reveal several key details of Na^+^ transport (Figure 2). Importantly, different binding positions of sodium ions within the pore were not accompanied by structural changes along the selectivity filter nor the entirety of the channel. Structure alignment along Cα resulted in a < 0.4 Å RMSD indicating high structural consistency. Previous crystal structures of NaK also observed no structural rearrangements when titrated with different Na/K ratios^35^ and in the presence of other monovalent (Rb^+^) and divalent cations (Ca^2+^, Ba^2+^)^11^. In contrast, computational electrophysiology, solution-state NMR studies and simulations demonstrated that the NaK SF can be structurally plastic and asymmetric, exhibiting several distinct conformations under different ions and salt concentrations^13,21,36–38^. SF asymmetry in an *I*4 space group may not be visualized because only one of the four polypeptides were found within the asymmetric unit while the other three are generated from crystal symmetry to form the tetramer. To our knowledge, other NaK SF conformations have not been captured experimentally.

For NaK2CNG, substitution of the SF to TVGDTPP mimics that of eukaryotic cyclic-nucleotide-gated (CNG) channels^15^. The SF consists of three contiguous sites (S2-S4) similar to K^+^ channels (Figure 2D). We determined 4 crystal structures of NaK2CNG purified in 150 mM KCl. Similar to NaK, after multiple rounds of refinement, different lateral positions of K^+^ along the conduction pore were resolved, depicting 8 distinct substates (Figure 2E). Alignment of all structures reveal less than 0.3 Å RMSD for all Cα hence, no structural changes were observed for the channel. Remarkably, each dataset processed revealed distinct structural features that are not universal to all. We observed two crystal substates showing the process of K^+^ dehydration as the ion approaches the SF (Supplementary Figure 1). Other structures of ion channels have shown hydrated ions in the external and the constriction sites, the locations of that channel that form the basis of permeation mechanisms^5,23^. This highlights the significance of sampling sufficient individual crystals to provide a more complete picture of the facets of the protein structure and dynamics.

### Ion-pore interactions

Three sites within the NaK SF were found with more preferential occupancy by Na^+^ - the external site (Sext), site 3 (S3) and constriction site (Scont) (Figure 3A). Notably, S3 had higher occupancy over other sites. We speculate that Sext and Scont are easily accessible for Na^+^ ions as they are close to external regions of the pore, while SV and S4 are most likely transient sites. In contrast, S3 likely provides a thermodynamically stable binding site for Na^+^ during ion conduction. The high occurrence of binding at S3 in our structures strongly agrees with earlier MD simulations that show Na^+^ ions can be tightly bound in-plane with the carbonyl oxygens of V64 and T63^23^. Consistent with MD simulations, we observed that not all Na^+^ ions are perfectly centered in S3 but rather exhibit three types of Na^+^-oxygen binding coordination: planar with V64 (configuration 1), square antiprism with both V64 and T63 (configuration 2), and planar with T63 (configuration 3, Figure 3C). MD simulations show Na^+^ ions fluctuate extensively near the upper and lower edges of the eight-carbonyl cage which can be experimentally recapitulated in our studies (Fig. 1, 3)^22,23,39^. The same simulation suggested that S3 imposes a discriminatory energy barrier for Na^+^, albeit low in energy^22^.

**Figure 3.**
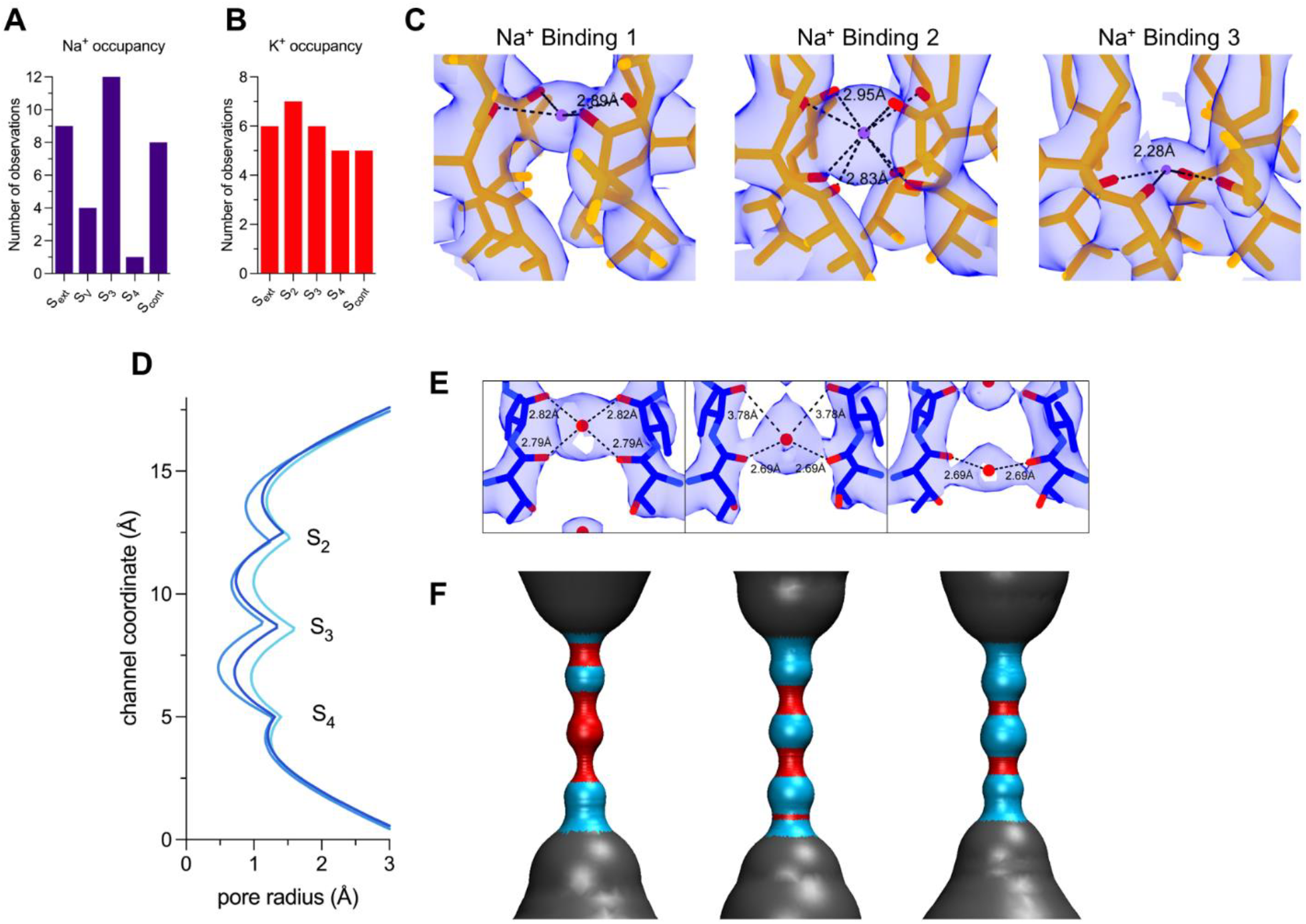
Interaction of Na^+^ and K^+^ with the selectivity filter. Tally of ions found in NaK (A) and NaK2CNG (B). (C) Electron density maps of the three types of binding configuration of Na in S3: planar with V64 carbonyls, square antiprism with carbonyls of T63, V64 and planar with T63 carbonyls. The distances between Na^+^ (purple) and the oxygens (red) are shown. (E) Density maps of the coordination of K^+^ (red) in S2/S3 and their corresponding HOLE profile (D,F). Opposite subunits are shown for ease.

While we didn’t observe water molecules within the NaK SF, it is possible that partial hydration occurs as the ions pass from the water-filled vestibule (SV) to S3. In this favorable planar configuration, ion-oxygen distances and partial exposure to water molecules provides Na^+^ a coordination number of 4-6, consistent with common Na^+^ coordination observed in water, proteins and other molecules^40^. In contrast, previous NaK structures only captured sodium ions situated perfectly in the center of each binding site, interacting in an eight-carbonyl cage similar to configuration 2^11,35,41^. Other NaK structures were not able to unambiguously assign positions for Na^+^ along the edges of S3/S411. Although ion occupancy in crystal structures is not a direct measurement of ion binding affinities, constant occurrence of binding in crystal structures statistically suggest the presence of a high affinity binding site. Consistent Na^+^ occupancy at S3 in all NaK structures suggest a binding mechanism more complicated than passive diffusion of ion.

Maps for K^+^ within the SF of NaK2CNG show no preference towards a particular binding site, in contrast to S3 in NaK (Figure 3B). This suggests a more concerted and transient movement of K^+^ down the conduction pore, most likely multiple ions at a time. Previous X-ray structures showed that the channel retains high K^+^ binding sites at S2 and S3 even at extremely low K concentration ratio (1 mM K^+^ / 49 mM Na^+^)^35^. The extra binding site (S2) replacing the vestibule (SV) in NaK would explain multiple binding of K^+^ but not why we observe the lack of a single high affinity binding site. Analysis of crystallographic data of K-selective channels shows that K^+^ ions form close-ion pairs at neighboring binding sites^9,20,42–44^. Previous molecular dynamics simulations suggested a strong ionic repulsion between adjacent K^+^ ions as they transverse the SF, a mechanism that could explain the high conduction rate of K^+^ ions in NaK2CNG while excluding water molecules^9^.

### K^+^-induced selectivity filter dilation

Alignment of all NaK and NaK2CNG structures indicated that no significant structural changes were induced by the different ion occupancy along the SF of the channels. To further investigate, the pore radius along the conduction pathway of the two tetrameric proteins were calculated using the program HOLE ^45^. For NaK, pore calculations revealed a relatively static pore profile, with no widening nor shrinking of the conduction pathway formed by the SF (Figure 4A). This is likely due to the size of Na^+^ fitting snuggly at the carbonyl sites of the SF.

**Figure 4.**
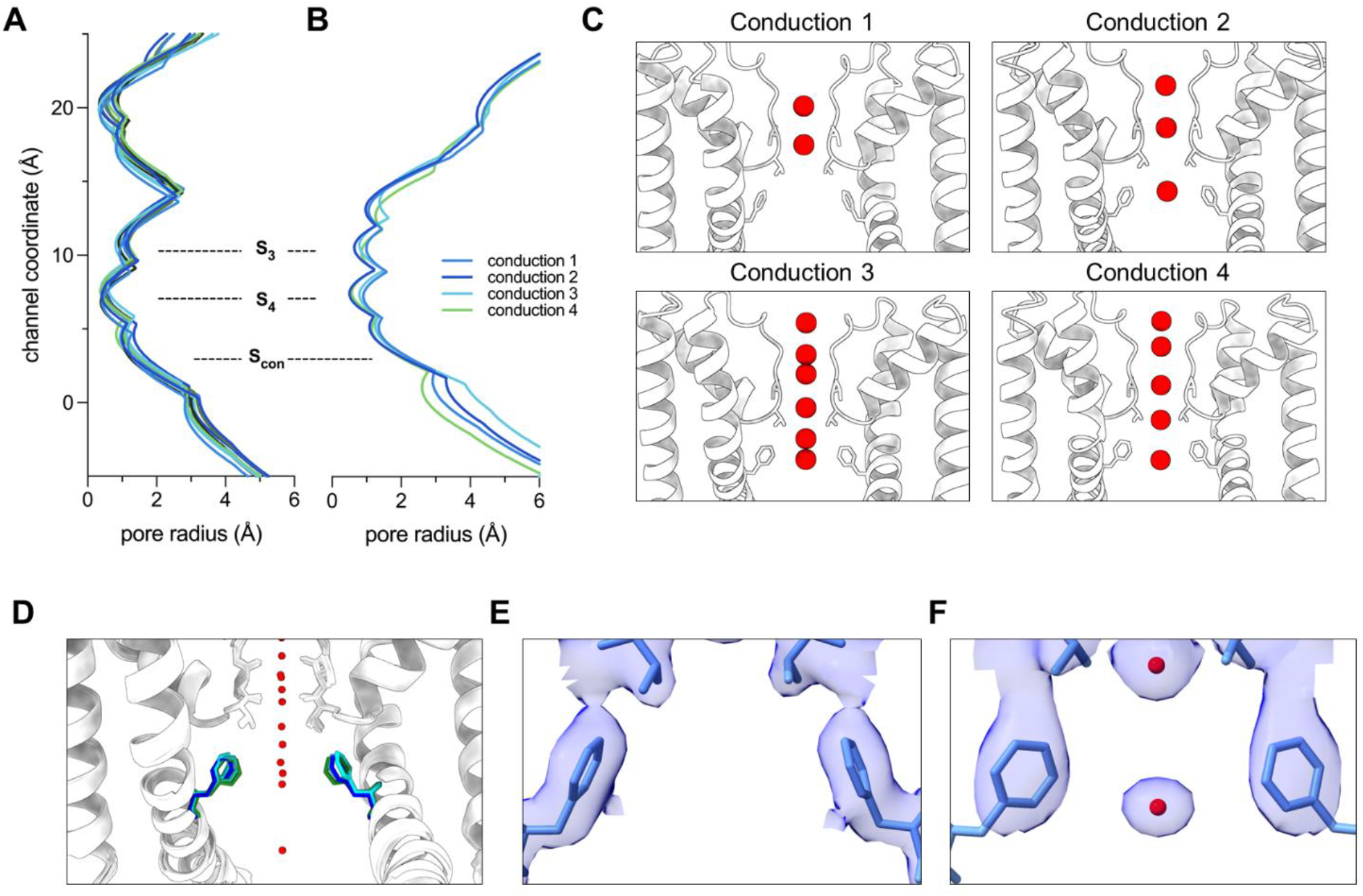
Gating at the restriction site. HOLE profiles of all NaK (A) and representative NaK2CNG (B), SF sites and constriction site are labelled. (C) K^+^ ions found within the conduction route of NaK2CNG with significant pore radius changes, labelled as conduction 1-4. (D) Alignment of four representative NaK2CNG highlighting F91. Electron density maps of F91 of conduction 1 (E) and conduction 4 (F).

In sharp contrast, the SF diameter varied significantly for NaK2CNG (Figure 3D). The tightest radius for both NaK and NaK2CNG is the planar cage formed by the carbonyls of T63 between S3 and S4, for NaK this ranges from 0.37 Å - 0.50 Å compared to 0.50 Å - 0.96 Å in NaK2CNG. Analysis of the tightest (0.50 Å), moderate (0.72 Å) and loosest (0.96 Å) pore sizes reveal that dilation of the T63 C-O SF lining is the result of K^+^ occupancy. Particularly, the pore dilates by approximately 0.5 Å when K^+^ directly interacts with the carbonyl oxygens of T63 in S4 in an imperfect planar binding. As K^+^ disengages from the carbonyl of T63, the pore returns to its contracted state producing its normal narrower T63 site. In the contracted state, K^+^ ions are perfectly caged in an 8-coordinate shell which is the preferred coordination of K^+^ within the SF. The dilation of S4, which serves as the intracellular entry site for cations, suggest that a permissive binding site is required for non-selectivity in ion channels. In this case, pore enlargement is likely essential to accommodate the need of K^+^ ions to fulfill an eight-coordinate shell.

The coordination numbers within the SF, whether the ion is interacting with the oxygens donated by carbonyls or waters, are known to range from 4 to 6 and 8^46^. According to the coordination model for selectivity, fluctuations in coordination numbers suggest structural distortions within the SF and diminishes the cavity rigidity, effectively reducing ion channel selectivity^47^. Because NaK2CNG is non-selective, a flexible and plastic selectivity filter could underly the mechanism for conduction of ions with sub-Ångstrom size differences. This work and several others challenge the original mechanistic view of a rigid cavity that fits a permeant ion is required for substrate discrimination in selective channels^13,48^. In fact, the notion of a rigid SF has been cautioned against by an overwhelming number of computational studies^14,49–51^.

### M2 helix allosteric gating

The constriction site has previously been identified as a gating mechanism for ion conduction that primarily involves the steric hindrances caused by F92 and its surrounding residues^12^. According to previous electrophysiology measurements, wildtype NaK and NaKΔ18 displayed almost negligible conduction rates while substitution of F92 into alanine results in drastic gain-of-flux^12,15,41,52^. In another work, the crystal structure of F92A NaK results in a wider constriction pore from 6.5 to 10.5 Å^12^. Here, close examination of the conduction route for NaK2CNG reveals significant variability in pore sizes.

The MicroED structures, and subsequent HOLE calculations, reveal that NaK did not undergo any pore alterations through the conduction pathway (Figure 4A). We believe that the functional effects of F92 may only be observed when it is mutated to alanine. However, for NaK2CNG, there is evident resizing of the pore starting from F91 towards the M2 helix. Particularly, pore resizing is related to the number of ions found along the conduction pathway (Figure 4B). We labelled the structures based on the number of resolved K^+^ ions along the conduction route, from two (conduction 1), three (conduction 2) and 5 or more (conduction 3 and 4, Figure 4C). The lowest conduction state (conduction 1) almost precluded the restriction of F91, widening the pore area from 2.6 Å to 3.9 Å. Alignment of all F91 shows a rotation of the phenyl ring of F91, fully facing the conduction route drastically widening the restriction site. On the other hand, we did not observe any correlation between the SF dilation and M2 helix movements.

The structural data presented here clearly demonstrates a degree of transmembrane allostery between the SF and the M2 helix, acting as an intracellular gate, as suggested by previous works ^13,21^. This is similar to the inter-gate coupling in MthK and KcsA thought to be important for channel activation and inactivation^53–56^. The phenylalanine residue found in the M2 helix is generally conserved in NaK and other channels (Supplementary Figure 2). Large aromatic residues in other channels have been proposed to act as a gate, such as Y132 in KirBac3.1, F145 in KirBac 1.1, F392/F434 in CNGA3/CNGB3 and F103 in KcsA^57–60^. In MthK, F92 residue position is equivalent to a small side chain amino acid (A88) which explains the lower conduction rates of NaK compared to MthK^61^. Previously published data suggested observable conformational flexibility that extends to the M2 helix but not big enough for a large-scale conformational change. Moreover, more recent NMR studies highlighted the functional effects of F92 where chemical shift perturbations were observed in the SF in F92A-mutated NaK, indicating conformational changes occur in the SF as a result of changes in F92 conduction pathway^52^.

The study presented here, along with previous *in silico* studies, suggest a more complicated channel gating than the opening and closing of the helical bundle crossing. The movements of the constriction site prime the residues in the SF for ion binding, ushering them through the pore. We speculate that the NaK channels are more actively participating in the ion transport than previously thought, although this requires further investigations.

## Conclusions

Here, several structures for NaK and NaK2CNG were sampled to probe channel dynamics which indicated both consistencies and subtle changes in SF conformation, in agreement with earlier biochemical and computational data. Since MicroED delivers charge density/Coulomb potential maps, this approach allowed us to unambiguously identify ions in the SF of the channels^30^. Through NaK, we were able to clearly locate Na^+^ positions along the selectivity filter that support theories in coordination. Moreover, since several structures were determined independently and compared, preferential binding of Na^+^ at several sites was identified. While these analyses did not identify structural changes along the pore of Nak, this was not the case for its mutant, NaK2CNG which did display several changes.

Despite significant SF homology to highly selective K^+^ channels, NaK2CNG remains indiscriminate towards monovalent cations. The data showed SF flexibility and dynamics. Specifically, we visualized two important ion channel properties, the K^+^-induced SF dilation and the M2 helix allosteric gating. Resolving ions along the SF allowed demonstration that the SF dilates to cater the coordination requirements of K^+^ ions. In part, the contraction of the S3 upon full coordination K^+^ can be observed as a stabilization effect by K^+^ ions. We believe that SF dilation principally rationalizes the non-selectivity of NaK2CNG despite similarities with K^+^-selective channels.

Multiple structures of NaK and NaK2CNG allowed us to also visualize the role of the constriction site in ion conduction. While NaK remained virtually static, NaK2CNG showed that the steric hindrance of F91 is nearly precluded at the low conduction state, opening the M2 helix. We speculate that this represents some of the initial steps of ion conduction. F91 opens as the first ions bind to the SF then closes as the ions exit towards the extracellular space. Moreover, our data strongly resembles the established C-type inactivation mechanism of the KcsA channel particularly the open-inactivated conformation where gate movements trigger conformational changes to the selectivity filter leading to a non-conductive state^59,62–64^. Conformational flexibility of the SF and M2 helix has previously been reported through solid state NMR and computational methods but has not been captured in crystal structures^13,21–24,36^.

MicroED has been at the forefront of structural biology of difficult targets, but it can also be used effectively to prove dynamics in protein samples even in the context of a nanocrystal. Indeed, through the NaK proteins, the high throughput method is effective and beneficial. Expansion of this work to other ion channels, such as highly selective ones, could offer additional valuable insights to the concepts of ion discrimination and permeation. We predict that as high throughput MicroED pipelines become more established and widespread additional studies would benefit from probing dynamics en route for a more complete biophysical representation of various mechanisms.

## Acknowledgements

This study was supported by the National Institutes of Health P41GM136508. Portions of this research or manuscript completion were developed with funding from the Department of Defense MCDC-2202-002. Effort sponsored by the U.S. Government under Other Transaction number W15QKN-16-9-1002 between the MCDC, and the Government. The US Government is authorized to reproduce and distribute reprints for Governmental purposes, notwithstanding any copyright notation thereon. The views and conclusions contained herein are those of the authors and should not be interpreted as necessarily representing the official policies or endorsements, either expressed or implied, of the U.S. Government. The PAH shall flow down these requirements to its sub awardees, at all tiers. The Gonen laboratory is supported by funds from the Howard Hughes Medical Institute.

## Accession Codes

Coordinates and maps were deposited in the protein data bank (Accession code XXXX) and the EM Data bank (Accession code YYYY).

## Methods

### Purification and Crystallization

The gene encoding for NaK and NaK2CNG, lacking the first 19 residues, were cloned into the pQE60 vector and transformed into BL21 cells. Preculture cells were grown at 37°C overnight in LB media with 100 µg/mL ampicillin. Each 1L of media were inoculated with 10 mL of preculture cells and grown for 4 hours at 37°C. Protein expression was induced by adding 0.4 mM isopropyl β-D-1-thiogalactopyranoside (IPTG) and incubating at 25°C for 20 hours. The cells were harvested by centrifugation at 4000 rpm using a JLA-8.1 rotor (Beckman Coulter). The resulting pellet was resuspended in a buffer containing 50 mM Tris (pH 8.0), 150 mM NaCl, 2 μg/ml DNase, 10 μg/ml lysozyme, and protease Inhibitors. The cell suspension was processed through a Microfluidizer (Microfluidics Corporation) at 15,000 psi, followed by ultracentrifugation at 42,000 rpm using a Ti45 rotor in an Optima L-90K Ultracentrifuge (Beckman Coulter) for 1 hour. The membrane pellet was resuspended in 50 mM Tris (pH 8.0) and 150 mM NaCl, and solubilized with 2% n-Decyl-β-D-maltoside (DM) at room temperature for 2 hours. Insoluble materials were removed by another round of ultracentrifugation at 42,000 rpm using a Ti70 rotor (Beckman Coulter) for 30 minutes. The supernatant was mixed with Talon beads pre-equilibrated in 50 mM Tris (pH 8.0), 150 mM NaCl, and 0.2% DM. The column was washed with 15 mM imidazole and eluted with 300 mM imidazole in the same buffer. The NaK protein was then incubated with thrombin protease at 4°C overnight. The cleaved proteins were further purified using a Superdex 200 size exclusion column (GE Healthcare) equilibrated in 50 mM Tris (pH 8.0), 150 mM NaCl, and 0.2% DM. The same purification was followed for NaK2CNG except 150 mM NaCl was substituted with 150 mM KCl.

Purified proteins were concentrated to 10 mg/mL using 30-kDa Sartorius concentrators. A condition matrix was prepared in a 96-well plate with (±)-2-methyl-2,4-pentanediol (MPD) concentrations ranging from 50% to 80% and 100 mM Tris pH 8.0. 0.2 µL of protein and 0.2 µL of reservoir solution was mized using a Mosquito crystallization robot. The plates were incubated at room temperature where microcrystals began to appear within 24 hours.

### EM grids preparation

Crystallization of NaK in Tris pH 8.0 and 60-80% (±)-2-methyl-2,4-pentadiol (MPD) produces micrometer-sized plate-like crystals that are barely visible under an optical microscope. Traditional blotting conditions were enough to prepare thin vitrified cryo-EM samples. EM grids were prepared in a Leica GP2 plunge freezer set to 95% humidity and 20°C temperature. Several optimizations were performed to achieve thin vitreous ice. Briefly, the crystals were 10x diluted with mother reservoir and loaded on a previously glow-discharged Quantifoil Cu 200 R2/2 (Quantifoil) holey carbon grids. The grids were blotted for 30s and plunge-frozen in liquid ethane.

### MicroED data collection

Clipped grids are loaded into a Titan Krios G3i TEM (Thermo Fisher) for data collection operating at an accelerating voltage of 300 kV. Grid atlases were acquired at 155x magnification with SerialEM to identify squares in thin ice containing possible crystals. Multiple good squares were selected using the “autocontour grid squares” function in SerialEM. Images of each square at 2250x magnification were taken. Crystals are visually identified wherein points are added within the crystal area. At each point, a single diffraction pattern was taken to segregate diffracting and non-diffracting crystals. All non-diffracting points were removed. Continuous rotation MicroED data were collected with a dedicated in-house SerialEM script on a Falcon 4i direct electron detector in electron counting mode, utilizing the electron event representation format (.eer). Each crystal was examined with a single dataset collected over a 70-degree wedge for 420 seconds, with a nominal camera length of 2500 mm.

### MicroED data processing

Each MicroED dataset was converted to SMV format using mrc2smv software. The converted frames were indexed, integrated and scaled with XDS. Molecular replacement was performed in Phaser using the PDB entry 3E89 for NaK and 3k03 for NaK2CNG. Structure refinement and modelling was carried out with phenix.refine and coot until the lowest R values were obtained. Conduction profiles in the final structures were calculated using the HOLE program.

**Supplementary Figure 1.**
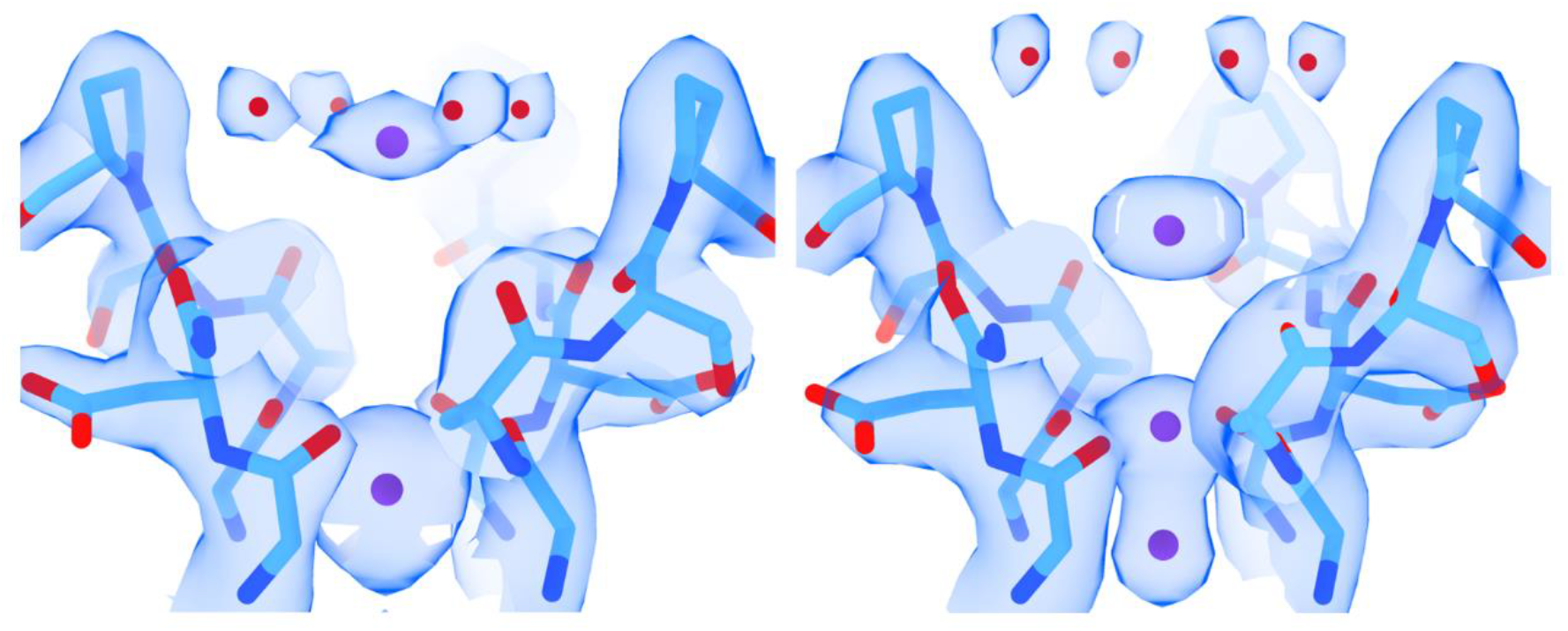
K^+^ dehydration along the NaK2CNG conduction pore.

**Supplementary Figure 2.**
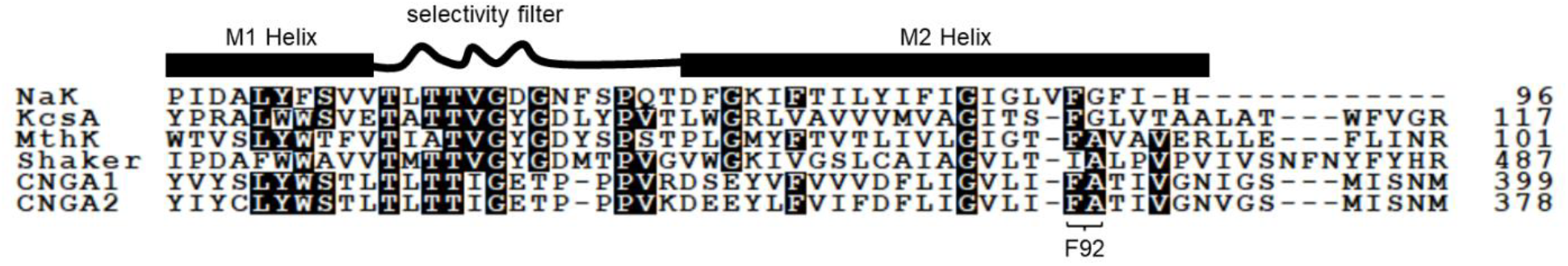
Sequence alignment of select ion channels.

